# Drosophilid cuticle pigmentation impacts body temperature

**DOI:** 10.1101/2022.12.03.518031

**Authors:** Laurent Freoa, Luis-Miguel Chevin, Philippe Christol, Sylvie Méléard, Michael Rera, Amandine Véber, Jean-Michel Gibert

## Abstract

Cuticle pigmentation has been clearly demonstrated to impact body temperature for several relatively large species of insects, but it was questioned for small insects. Here we used a thermal camera to assess the impact of drosophilid cuticle pigmentation on body temperature when individuals are exposed to light. We compared mutants of large effects within species (*Drosophila melanogaster ebony* and *yellow* mutants). Then we analyzed the impact of naturally occurring pigmentation variation within species complexes (*Drosophila americana/Drosophila novamexicana* and *Drosophila yakuba/Drosophila santomea*). Finally we analyzed lines of *D. melanogaster* with moderate differences in pigmentation. We found significant differences in temperatures for each of the four pairs we analyzed. The temperature differences appeared to be proportional to the differently pigmented area: between *Drosophila melanogaster ebony* and *yellow* mutants or between *Drosophila americana* and *Drosophila novamexicana*, for which the whole body is differently pigmented, the difference in temperatures was around 0.6°C ±0.2°C. By contrast, between *D. yakuba* and *D. santomea* or between *Drosophila melanogaster Dark* and *Pale* lines, for which only the posterior abdomen is differentially pigmented, we detected a temperature difference of about 0.14°C ±0.10°C. This demonstrates that cuticle pigmentation has ecological implications in drosophilids regarding adaptation to environmental temperature.

## Introduction

Drosophilid pigmentation has been used as a fruitful model to dissect the molecular bases of sexual dimorphism and morphological variation and evolution ^1–4^. Indeed, it is a particularly rapidly evolving trait, such that different populations or closely related species can have dramatically different pigmentations ^5–7^. In contrast, the ecological relevance of pigmentation is much less well known, and its effects on fitness are difficult to establish in the field, as this trait is pleiotropically linked to many other traits affecting fitness, such as life history (longevity, fecundity), cuticular hydrocarbons, and resistance against pathogens, parasites, UV or desiccation ^8–14^. The direct influence of pigmentation, independent from other traits to which it may be correlated in the field, can instead be assessed by measuring its effect on aspects of performance (sensu Arnold 1983^15^) related to specific hypotheses, in controlled environments. For instance, a common hypothesis is that drosophilid pigmentation plays a role in thermoregulation, and thus in their adaptation to environmental temperature ^16^. Dark-colored flies may warm up more in the sun, while light-colored flies may avoid overheating. In agreement with this hypothesis, in *Drosophila melanogaster*, populations living at higher altitudes or higher latitudes are darker ^5,17–20^ and abdominal pigmentation shows some phenotypic plasticity ^16^: flies which develop at low temperature are darker, which is thought to be adaptive. The influence of pigmentation on body temperature was shown in many ectotherms (thermal melanism) ^21^ and even in distantly related organisms such as yeasts ^22^. In insects, it has been demonstrated in species from several orders (Orthoptera, Hemiptera, Coleoptera, Lepidoptera) ^23–28^. However, all these insect species have relatively large sizes. It was shown, using pairs of insects of comparable sizes and different pigmentations, that the effect of pigmentation on the temperature of insects exposed to sunlight was clear for large insects but was extremely limited for small insects (around 3mg) ^29^. For such small body sizes and with the calorimetry tools available at the time, it was not possible to conclude on the existence of a relation between body temperature and pigmentation ^29^. Drosophilids usually have a smaller weight (between 1 and 1.5 mg fresh weight for a *Drosophila melanogaster* female ^30^) than the insects used in this previous study ^29^, which makes the impact of drosophilid pigmentation on body temperature unclear. In this work, we used a thermal camera equipped with a macro lens to monitor the body temperature of drosophilids exposed to a light source mimicking sunlight, to assess the role of pigmentation on body temperature in these organisms. Thermoregulation was treated as an element of performance affected by pigmentation, and thus as a proxy for fitness. We tested pairs of Drosophila lines or species differing by their pigmentations over their whole body, or only over some portion of their abdomens. These differences in pigmentation have been previously described and their genetic bases characterized ^6,7,31–36^. The choice of these pairs of lines or species was based on the existence of strong phenotypic differences within the same species (*Drosophila melanogaster ebony* and *yellow* mutants), natural genetic variation within the same species (*Drosophila melanogaster Dark* and *Pale* lines), or different pigmentation in very closely related species with otherwise similar morphology (*Drosophila americana/Drosophila novamexicana* and *Drosophila yakuba/Drosophila santomea*). We compared the evolution of body temperature between the darkest fly and the lightest fly using the thermal camera, which allowed us to visualize very small differences in temperature (as low as 0.05°C).

## Results

We divided the results into five sections. The first section compares mutants of large effects within species (*Drosophila melanogaster ebony* and *yellow* mutants). The second and the third sections concern naturally occurring variation within species complexes (suggestive of local adaptation), with either whole-body or anatomically restricted pigmentation differences (respectively *Drosophila americana/Drosophila novamexicana* and *Drosophila yakuba/Drosophila santomea*). The fourth section focuses on lines of *D. melanogaster* obtained by artificial selection with moderate differences in pigmentation. In each section, we give detailed information on the lines or species used. Temperature measures are available in Tables S1-S8 (see Material and Methods for their treatment). The fifth section analyses the relationship between pigmentation difference and body temperature difference.

### *ebony* and *yellow Drosophila melanogaster*

In order to compare *Drosophila melanogaster* individuals with very different pigmentations, we used loss of function alleles of *ebony* (*ebony^1^, e^1^*) and *yellow* (*yellow^1^, y^1^*). The *e^1^* allele blocks the production of yellow NßAD sclerotin (Figure 1), such that the fly cuticle is strongly melanized as more dopamine is available to produce black and brown melanins. Conversely, *y^1^* flies cannot produce black melanin (Figure 1) and their cuticle is pigmented only with brown melanin and yellow NßAD sclerotin.

**Figure 1:**
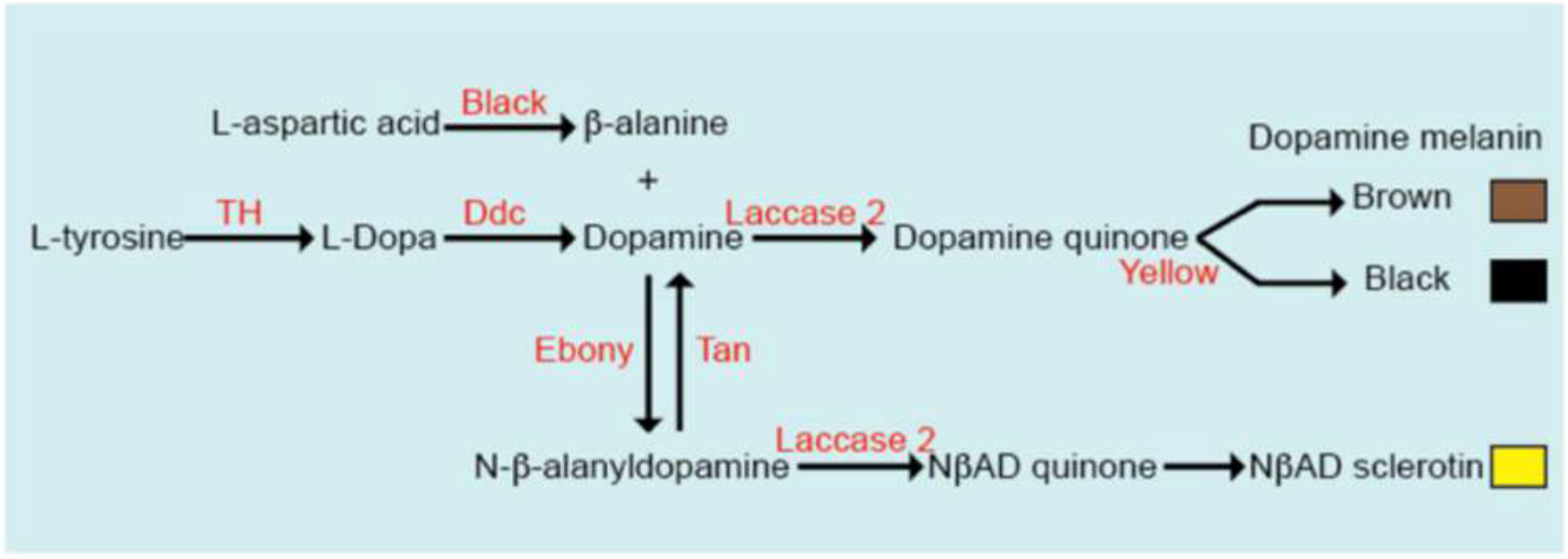
Synthesis pathway of cuticle pigments in *Drosophila melanogaster*.

Thus, despite belonging to the same species, *e^1^* and *y^1^* flies have dramatically different pigmentations (see Figure 2a).

**Figure 2:**
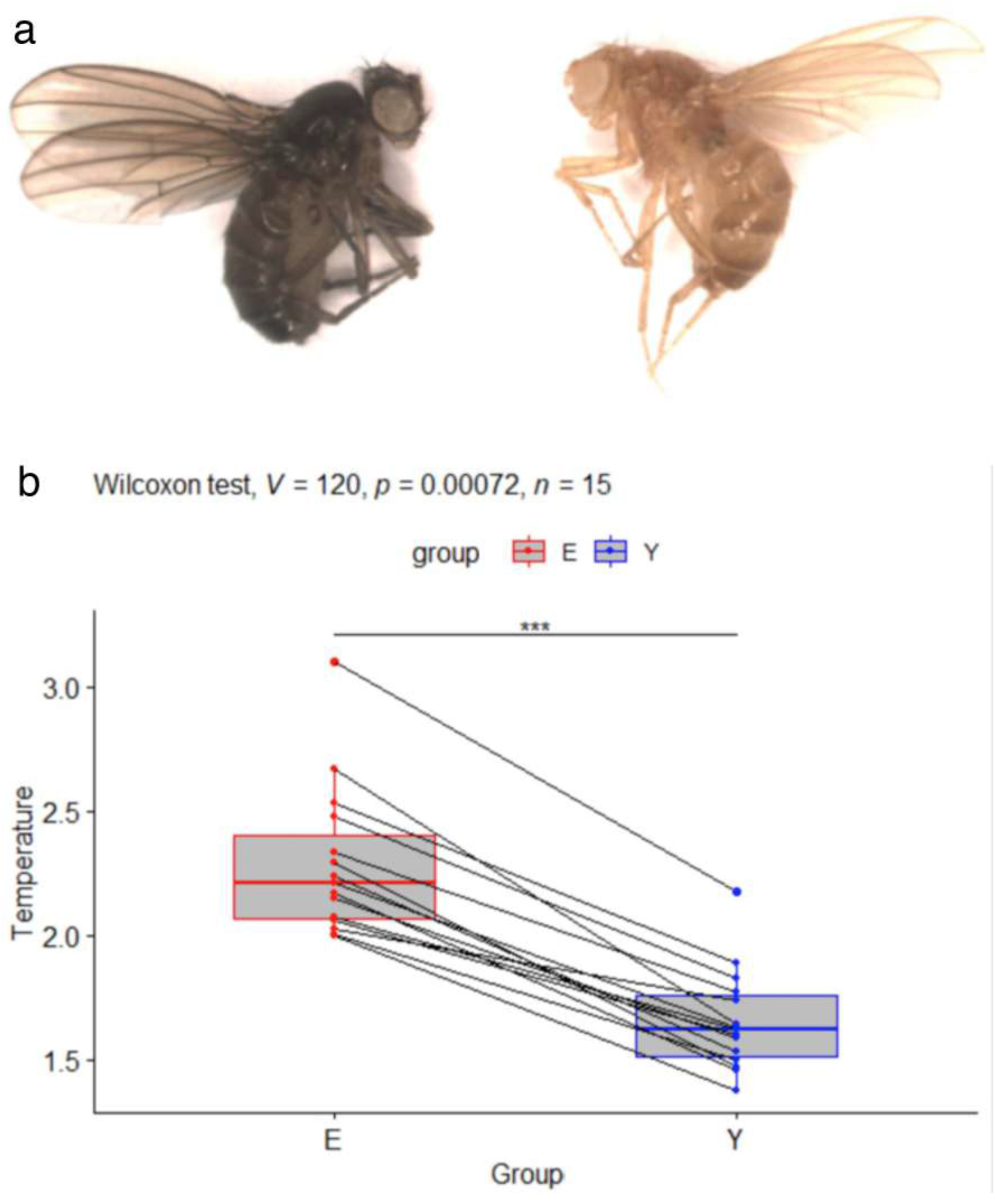
a: Picture of *ebony^1^* (left) and *yellow^1^* (right) *Drosophila melanogaster* females. b: Boxplots showing the normalized temperatures in °C for *D. melanogaster ebony (E*) and *yellow (Y*) mutant females. Pairs of individuals recorded simultaneously are indicated by lines. In all pairs, the *ebony* fly is hotter than the *yellow* fly. This is confirmed by a Wilcoxon rank signed test showing that E is significantly hotter than Y (p-value p = 0.00072, V = 120 being the value of the test statistic). ***: p<0.001

A Wilcoxon signed rank test on paired samples was performed on the subset of data_norm corresponding to pairs of *ebony^1^* and *yellow^1^* flies to test whether flies with different genotypes (leading to different pigmentations) had different temperatures when exposed to light. This test revealed a significant effect of the genotype on the temperature of the flies (p<0.001), allowing us to conclude that the difference of pigmentation between *ebony* and *yellow* flies indeed impacted their body temperature when lit up with sun-mimicking lighting.

Figure 2b shows significant variation between the 15 experimental replicates. However, in each replicate, the *y^1^* fly was constantly colder than the *e^1^* fly. The average temperature difference between *e^1^* and *y^1^* females was 0.63±0.19°C.

### *Drosophila americana* and *Drosophila novamexicana*

*Drosophila americana* and *Drosophila novamexicana* are sister species within the *Drosophila virilis* species group and diverged recently, about 300,000 to 500,000 years ago ^6^. The body colour of *Drosophila novamexicana* has a derived yellow pigmentation, while the colour of other members of this group (including *Drosophila americana*) is dark brown ^6^ (see Figure 3a).

**Figure 3:**
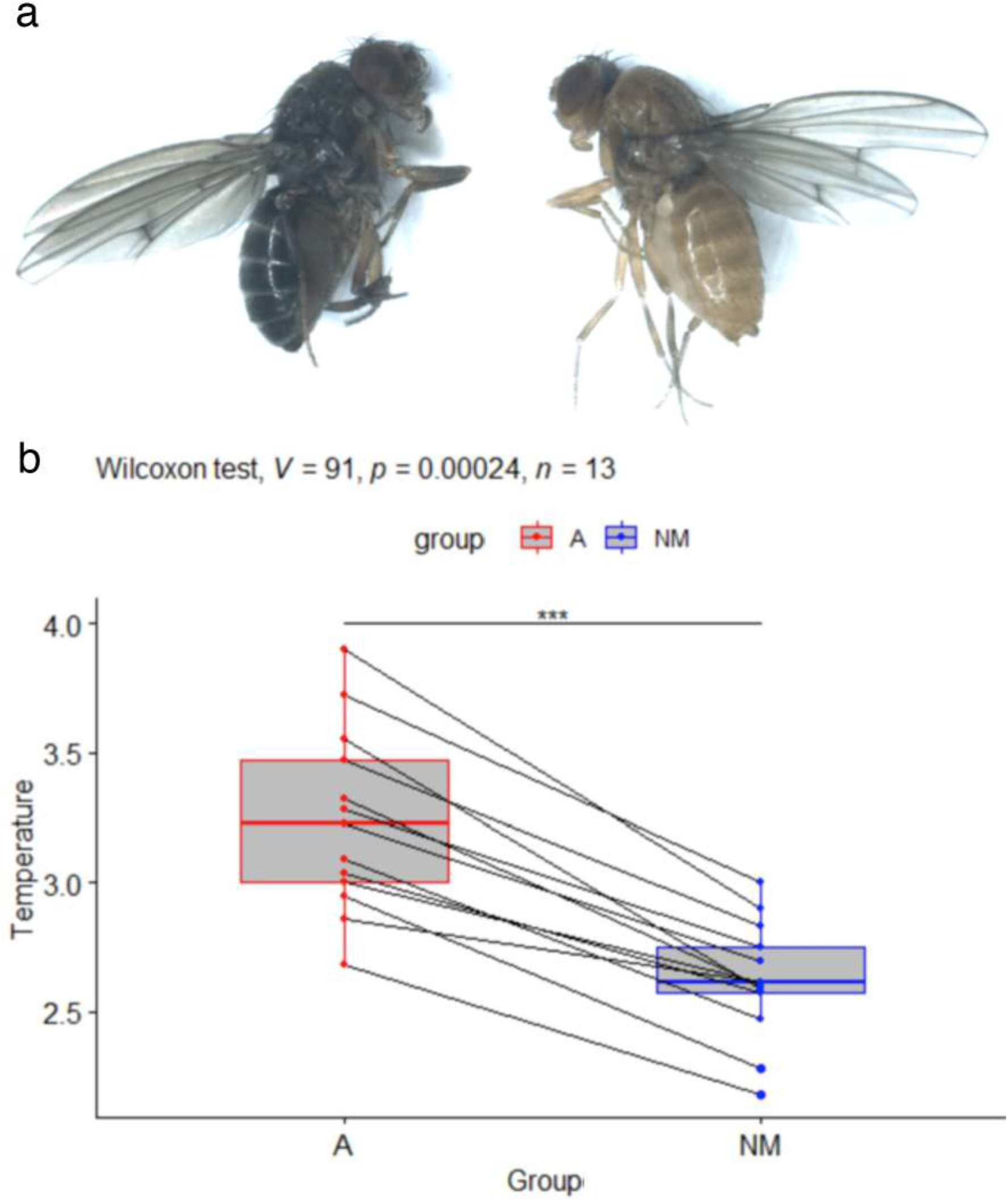
a: Picture of *D. americana* (left) and *D. novamexicana* (right). b: Boxplots showing the normalized temperatures in °C for *D. americana* (A) and *D. novamexicana* (NM) females. Pairs of individuals recorded simultaneously are indicated by lines. In all pairs, the *D. americana* individual is hotter than the *D. novamexicana* individual. This is confirmed by a Wilcoxon rank signed test showing that A is significantly hotter than NM (p-value p = 0.00024, V = 91 being the value of the test statistic). ***: p<0.001

These species are native to North America. *D. novamexicana* is localized in the arid south-western regions of the USA and Mexico, whereas *D. americana* extends over a wide geographical and climatic range, from the western Great Plains to the east coast of North America ^37^. The occurrence of *D. novamexicana* in an arid zone at one edge of the range of *D. americana* and its lighter pigmentation suggests that this species is specialized to this hotter habitat. In the laboratory, these species can mate and produce fertile offspring. Genetic mapping has shown that genomic regions containing the *ebony* and *tan* genes contributed to the pigmentation divergence between *D. novamexicana* and *D. americana*^6^ and further studies confirmed the role of both genes ^32,33^.

We observed that body temperatures were significantly different between the 2 species (p<0.001, see Figure 3b). There is again a strong variation between replicates, but in each replicate the body temperature of the *D. novamexicana* fly was always lower than that of the *D. americana* fly (Figure 3b). The average temperature difference between *D. americana* and *D. novamexicana* females was 0.61±0.21°C.

### *Drosophila yakuba* and *Drosophila santomea*

This pair of closely related species belongs to the *Drosophila melanogaster* species group. They diverged between 500 000 years and 1 million years ago ^38^. *Drosophila yakuba* is widely present on the African continent and on several African islands, whereas *Drosophila santomea* is endemic of the Island of Sao Tome, where it co-occurs with *Drosophila yakuba*^39^. They show contrasting pigmentation patterns: in both sexes, *Drosophila santomea* has a pure yellow body color, without the black pattern observed in *Drosophila yakuba* and other species of the *Drosophila melanogaster* group ^39^. These two species have a reduced sexual dimorphism compared to the pigmentation of other species of the *Drosophila melanogaster* subgroup, where the last segments of the abdomen of females are less pigmented than those of males. The difference in pigmentation between these two species is however maximal in males, in which abdominal segments 5 and 6 are fully melanized in *Drosophila yakuba*, but homogeneously yellow in *Drosophila santomea* (see Figure 4a). The difference in pigmentation between these species is much more localized than between the previous species.

**Figure 4:**
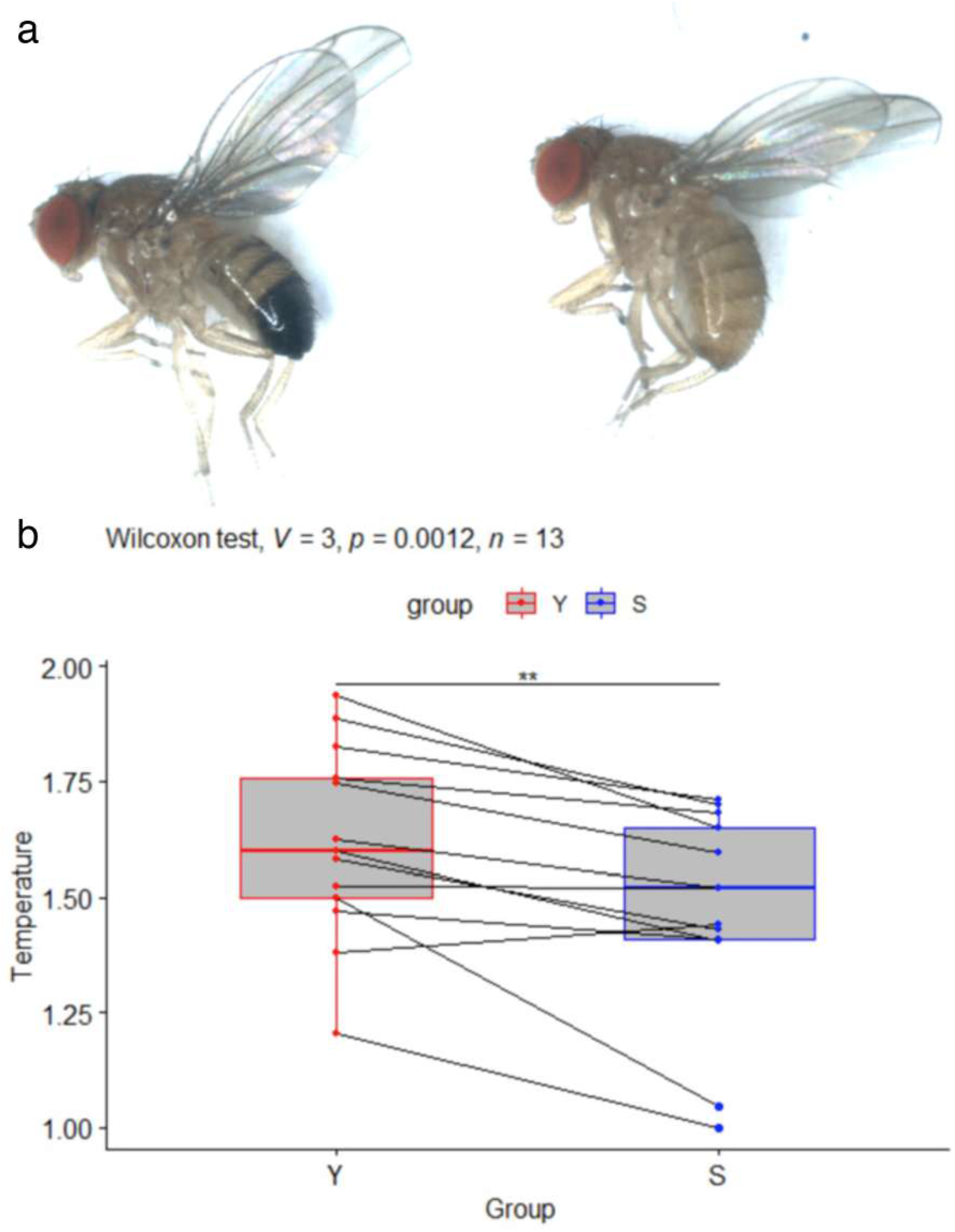
a: Picture of *Drosophila yakuba* (left) and *Drosophila santomea* (right) males. b: Boxplots showing the normalized temperatures in °C for *D. yakuba* (Y) and *D*. *santomea* (S) males. Pairs of individuals recorded simultaneously are indicated by lines. In all pairs but one, the *D. yakuba* individual is hotter than the *D. santomea* individual. This is confirmed by a Wilcoxon rank signed test showing that Y is significantly hotter than S (p-value p = 0.0012, V= 3 being the value of the test statistic). **: p<0.01

In the laboratory, these species can mate and produce fertile hybrid females, but sterile males (consistent with the classical pattern described as Haldane’s rule ^40^). There is evidence from field studies and population genetics that hybridization occurs in the wild between these species on the island of Sao Tome ^41^. Genetic analyzes indicated that at least 5 loci are responsible for the difference in pigmentation between *D.yakuba* and *D. santomea:* the pigmentation enzyme coding genes *yellow* (*y*), *tan* (*t*) and *ebony* (*e*) and the genes encoding the transcription factors Abdominal-B (*Abd-B*) and Pdm3 (*pdm3*)^34^. A recent study based on artificial introgression identified an additional locus involved, *Grunge* (*Gug*)^35^. Interestingly, long-term introgression experiments of pigmentation genes between *Drosophilia santomea* and *Drosophilia yakuba* revealed pigmentation-based assortative mating, ^35^ which suggests that pigmentation differences contribute to reproductive isolation between these species.

As for previous comparisons, we found that body temperatures were significantly different between the two species (p<0.01). Despite the variations between replicates, in all replicates but one, the *D. santomea* individual was observed to be colder than the *D. yakuba* individual (Figure 4b). The average temperature difference between *D. yakuba* and *D. santomea* males was 0.15±0.13°C.

### *Drosophila melanogaster Dark* and *Pale* lines

These two lines were generated by artificial selection starting from a *Drosophila melanogaster* Canadian population that was polymorphic for female abdominal pigmentation ^31^. Each line was isogenized through brother-sister crosses for 10 generations. The pigmentation difference between females of these two lines is located in the posterior abdomen (see Figure 5a) and is mainly caused by allelic variation at the *bric-à-brac* locus encoding the transcription factors *bab1* and *bab2*^31^. Indeed, in the enhancer driving *bab* gene expression in posterior abdominal epidermis, there is a deletion removing two Abdominal-B binding sites in the Dark line which reduces the activity of the enhancer ^31^.

**Figure 5:**
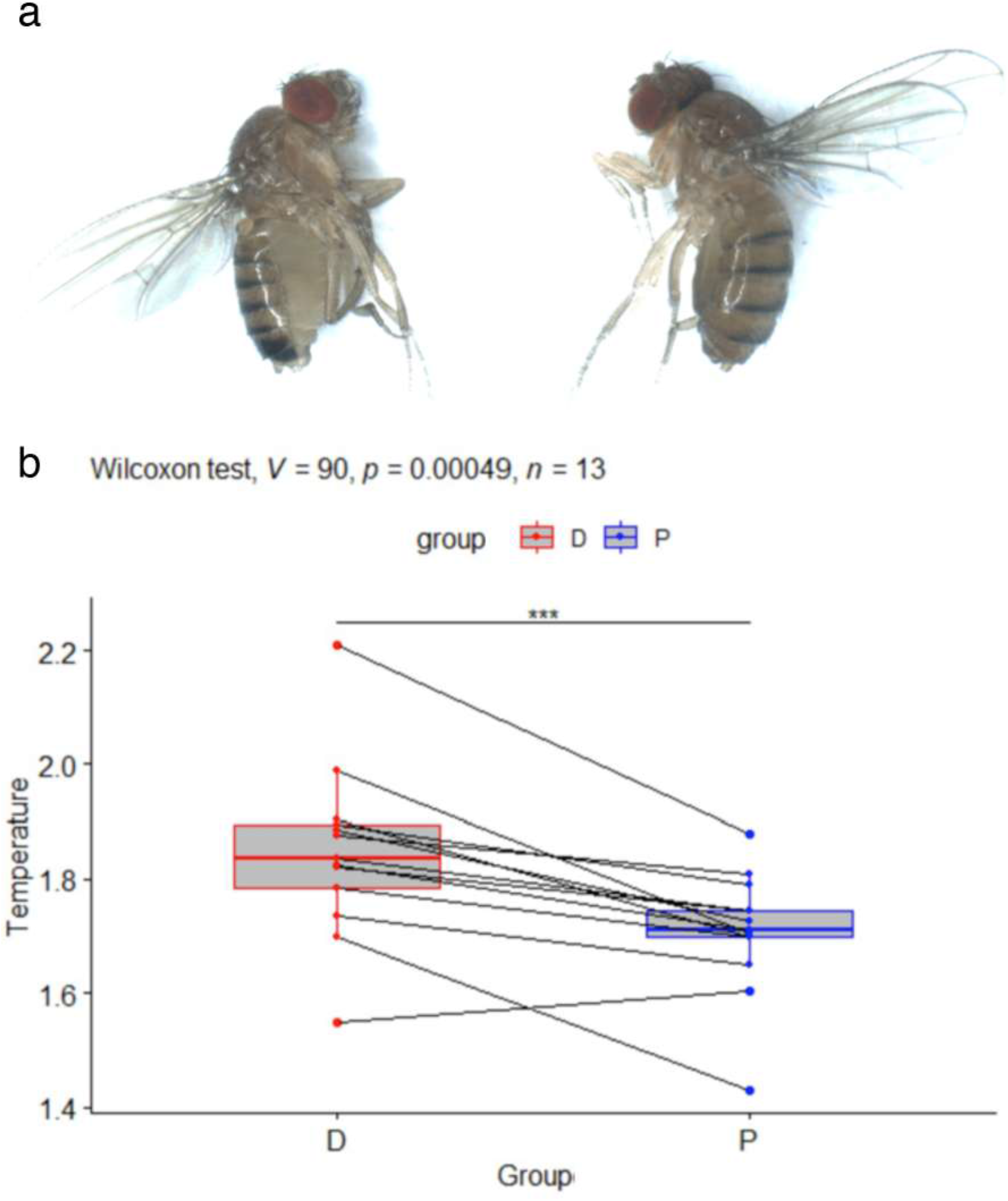
a: Picture of *Drosophila melanogaster Dark* (left) and *Pale* (right) females. b: Boxplots showing the normalized temperatures in °C for *D. melanogaster Dark* (D) and *Pale* (P) females. Pairs of individuals recorded simultaneously are indicated by lines. In all pairs but one, the *Dark* individual is hotter than the *Pale* individual. This is confirmed by a Wilcoxon rank signed test showing that D is significantly hotter than P (p-value p = 0.00049, with V = 90 being the value of the test statistic). ***:p<0.001

Again, body temperatures were significantly different between individuals of the 2 genotypes (p<0.001) despite strong variation between replicates. Indeed, in all replicates but one the *Pale* fly was observed to be colder than the *Dark* fly (Figure 5b). The average temperature difference between *D. melanogaster Dark* and *Pale* females was 0.14±0.11°C.

### The temperature difference is related to the difference in pigmentation

In order to visualize the relation between pigmentation differences and temperature differences for the four pairs of fly comparisons, we plotted them on the same graph. For this, we measured pigmentation differences of 10 pairs of flies for each of the four comparisons (thorax and abdomen, see Material and Methods): *ebony-yellow* (Table S9), *D. americana-D. novamexicana* (Table S10), *D. yakuba-D. santomea* (Table S11) and *D. melanogaster Dark-Pale* (Table S12). The graph shows that the difference in temperature is related to the difference in pigmentation (Figure 6).

**Figure 6:**
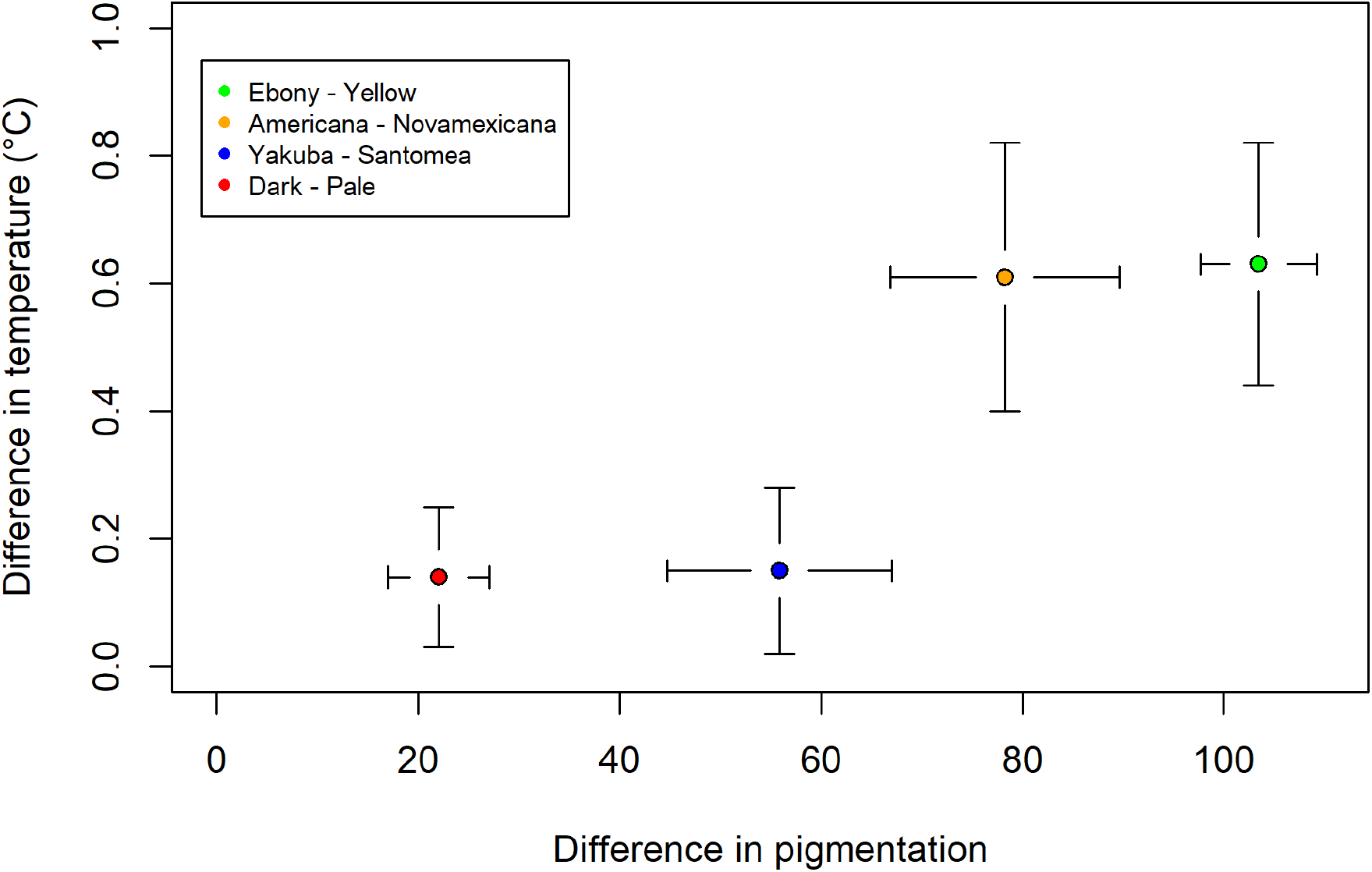
Graph showing the relation between pigmentation differences and temperature differences for the four pairs of comparisons (means and standard deviations). Pigmentation and temperature were not measured on the same individuals.

It is maximum (around 0.6°C) for the most differently pigmented flies (*ebony-yellow* and *D. americana-D. novamexicana*), for which the whole body is differently pigmented, and smaller (around 0.14°C) for the least differently pigmented ones (*D. yakuba-D. santomea* and *D. melanogaster Dark-Pale*) for which the pigmentation difference is localized to the posterior abdomen.

## Discussion

Here, we showed that for the 4 pairs of Drosophila species or lines that we compared, the most pigmented Drosophila in each pair was warmer than the less pigmented one when exposed to a light source mimicking sunlight. The temperature difference appeared to be proportional to the differently pigmented area: between *Drosophila melanogaster e^1^* and *y^1^* mutants or between *Drosophila americana* and *Drosophila novamexicana*, for which the whole body is differently pigmented, the difference in temperatures was approximately 0.6°C ±0.2°C. By contrast, between *D. yakuba* and *D. santomea* or between *Drosophila melanogaster Dark* and *Pale* lines, for which only the posterior abdomen is differentially pigmented, we detected a temperature difference of about 0.14°C ±0.10°C. Thus, although the impact of pigmentation on body temperature was previously undetected for small insects ^29^, using the thermal camera we could measure temperature differences between drosophilids of different pigmentation, even if they were of low magnitude. These effects of pigmentation on body temperature are likely to have ecological impacts. For example, the derived light pigmentation of *D. novamexicana*, which we showed to have an impact on body temperature, could have helped this species to adapt to the hot desert areas where it lives ^37^.

We showed that natural genetic variation for pigmentation within species had an effect on body temperature (*D. melanogaster Dark* and *Pale* line). For *D. yakuba, D. santomea, D. novamexicana* and *D. americana*, only one line per species was analyzed. However, in species such as *Drosophila americana* for example ^6,37^, it was shown that there was genetic variation for pigmentation. It would then be interesting to investigate how such variation affects body temperature.

Our results show that thermal melanism applies to drosophilids. Thus, we expect that drosophilid pigmentation should vary with spatial and temporal gradients of temperature that influence natural selection in the field. It is already known that populations of *D. melanogaster* living at high altitude in Africa and India are darker ^5,17^. Similarly, *D. melanogatser* thoracic pigmentation is darker at high latitudes ^18–20^. Furthermore, *D. melanogaster* developed at low temperature show a darker pigmentation, which is thought to be an adaptive trait ^16^. Thus, it would be interesting to elaborate a model showing how genetic variation for pigmentation is modulated by spatial and temporal variations of temperature. This model would take into account that pigmentation is modulated both by genetic variation and by the temperature at which development takes place. It was shown that there is latitudinal and seasonal genetic variation in *Drosophila melanogaster*^42–44^. However, it is not known whether this variation involves allele frequencies of genes involved in abdominal pigmentation, although there is latitudinal variation for thoracic pigmentation ^18–20^. A related and timely issue is whether global warming will affect the genetic variation for pigmentation in drosophilids, as it was shown to have an impact on the distribution of species of butterflies and dragonflies of particular pigmentation in Europe ^45^ and on pigmentation variation in ladybirds ^46^ and some species of leaf beetles ^47^. Indeed, it was already shown that global warming had a detectable impact on genetic variation in particular species of drosophilids ^48^.

Our demonstration that pigmentation affects body temperature in drosophilids opens the way for studies investigating the fitness consequences of this trait, and therefore how natural selection operates on it. In several insect species, the effect of pigmentation on body temperature has an impact on global activity ^24–26^. Thus, it would be interesting to test whether we can detect an effect of pigmentation in drosophilids on activity, for example by measuring locomotion performance. More generally, the impact of body temperature on life-history components of fitness (such as age at maturity or fertility) is important to understand how selection operates on traits affecting thermal regulation, such as pigmentation. This may also explain how and why anatomically localized pigmentation may be favored, if the temperature of some organs (such as gonads) is more determinant to fitness that others. For any such studies, the results we report here will provide a much-needed quantitative baseline for relating pigmentation to temperature, and thus connect to the abundant literature on thermal adaptation.

## Materials and methods

### Origin of the drosophilids

The *Drosophila melanogaster* alleles *ebony^1^ (e^1^*) and *yellow^1^ (y^1^*) were obtained from the Bloomington Drosophila Stock Center (Reference BL1658 and BL169). In order to be assessed in the same genetic background, they were introgressed for more than 8 generations in the *w^1118^* stock.

*Drosophila americana* (line *w11*) and *Drosophila novamexicana* (line *15010-1031-04*) were provided by Jorge Vieira (University of Porto, Portugal).

*Drosophila yakuba* was provided by the late Jean David (EGCE, Gif sur Yvette, France) and *Drosophila santomea* (line *Cago 315*) was provided by Virginie Courtier-Orgogozo (Institut Jacque Monod, Paris, France).

The *Drosophila melanogaster* lines *Dark* and *Pale* were generated by artificial selection starting from a population polymorphic for female abdominal pigmentation and were previously described ^31^.

Flies were grown on standard medium at 25°C.

### Infrared thermography experiments

A FLIR thermal camera (FLIR A655sc) equipped with a macro lens (FLIR 2.9x) was used to image flies in the infrared spectrum for a given time interval.

During the experiment, flies were exposed to a source of light mimicking sunlight (25w, Repti Basking Spot Lamp, ZOO MED Europe).

The infrared thermography experiments were performed in an incubator maintaining a temperature of 16°C (POL EKO ST3 BASIC SMART). This prevented temperature disturbances due to external events except the ignition of the lamp, and allowed the experiments to start at similar temperatures.

The software FLIR ResearchIR Max was used to acquire and treat infrared thermography images. We used the following parameters: Emissivity: 0.95; Distance: 0.1m; Reflected Temp: 20°C; Atmospheric Temp: 16°C; Relative humidity: 50%; Transmission: 1; External optic: 16°C; Transmission: 1.

Flies were anesthetized using vapors of flynap (50% triethylamine, 25% ethanol, 25% water). For each experiment, a dark-colored fly and a light-colored fly were filmed simultaneously and side by side with the thermal camera in order to minimize acquisition biases. During each recording, flies were placed in the incubator, on a white paper, close to each other and equidistant from the camera and the lamp. These positions were chosen for the flies to be subjected to the same influence of the lamp when it was switched on. The recording of the thermal camera began when the average surface temperature measured by the camera was close to 16°C. Each recording lasted 3min30s and contained 1245 images. Starting at timestamp 30 seconds after the beginning of the recording, we switched on the lamp until timestamp 2min30. The recording was stopped at 3min, giving access to the temperature decrease dynamics. At the end of the first recording, the position of the flies were reversed in order to minimize the potential non-homogeneity of the illumination of the lamp on the surface, thus preventing a position effect. The recording was then reproduced identically to the previous one in this new configuration. This experiment was repeated on several pairs of flies, and for several pairs of fly species or lines: we carried out 15 comparisons of *Drosophila melanogaster ebony* and *yellow* flies, 13 comparisons of *Drosophila americana* and *Drosophila novamexicana* flies, 13 comparisons of *Drosophila yakuba* and *Drosophila santomea* flies, and 13 comparisons of flies from the *Drosophila melanogaster Dark* and *Pale* lines.

We illustrate the type of data collected with the experiment on *ebony^1^* and *yellow^1^ D. melanogaster* mutants shown in Figure 7.

**Figure 7:**
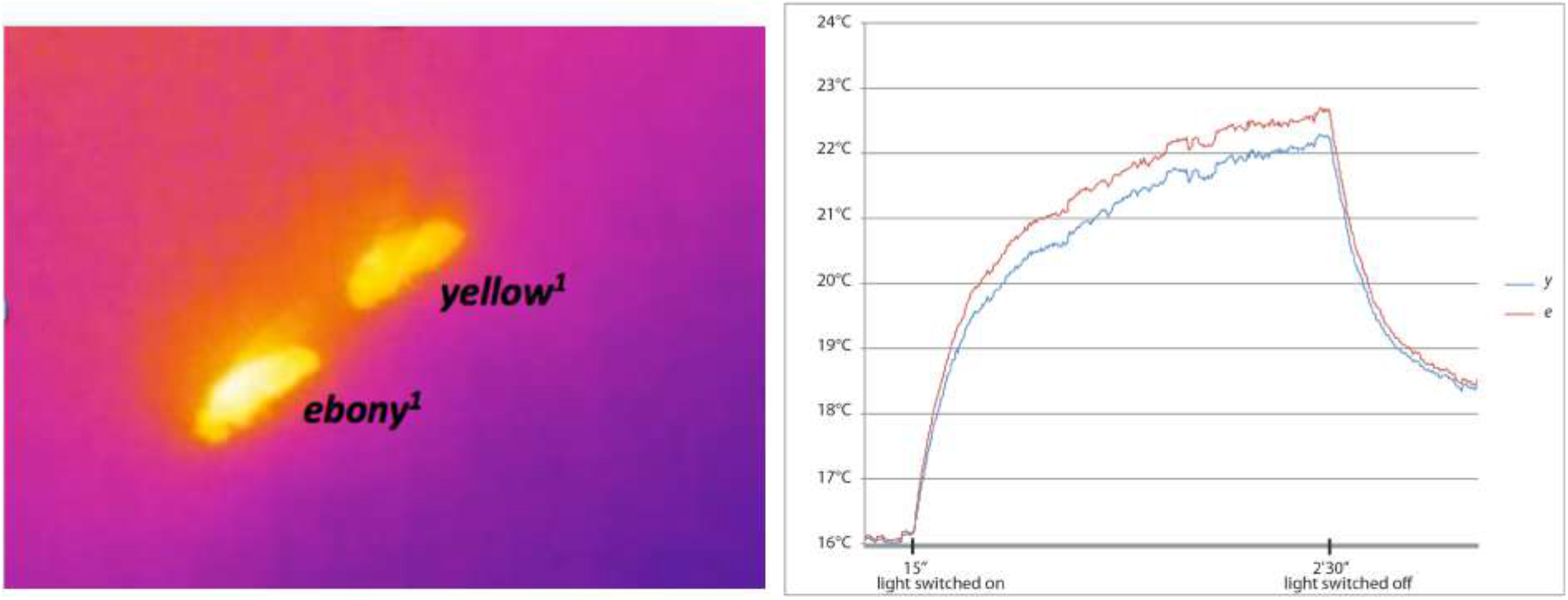
Evolution of fly body temperatures in an experiment with *D. melanogaster ebony* and *yellow* mutants. a: snapshot taken after the light was switched on (lowest temperature is blue, hottest temperature is white). b: temperature curves of the two flies recorded during the whole course of the experiment.

In this experiment, we see that the body temperature of the *ebony^1^* fly was observed to be higher than the temperature of the *yellow^1^* fly (see Figure 7a). When the lamp was switched on, the temperature of the two flies increased rapidly and a difference in temperatures between the two flies emerged after 15s (Figure 7b).

In order to further reduce any position effect and possible variations between experiments, we subtracted the average temperature of the paper surrounding each fly to the temperature of the fly. To do so, we drew ellipses in zones around each fly and called “temperature in the ellipse” the mean temperature in the area covered by the ellipse (see Figure 8).

**Figure 8:**
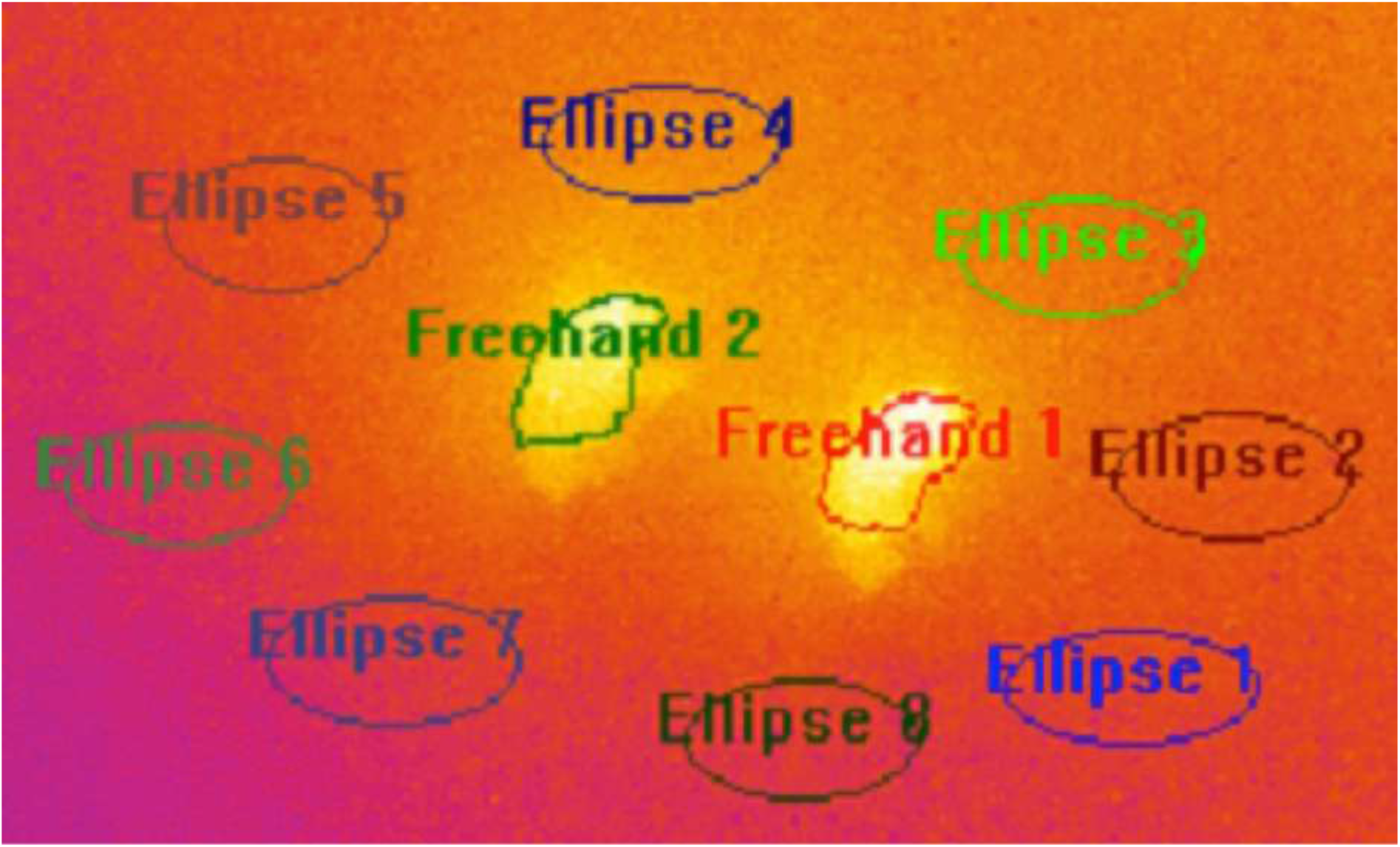
Principle of temperature normalization. In this screenshot of a video taken with the FLIR camera, *Drosophila santomea* is on the left and *Drosophila yakuba* is on the right. The flies are surrounded by eight ellipses numbered 1 to 8. The regions denoted by “Freehand” delimit the areas covered by the bodies of the two flies monitored during the experiment. In this example, ellipses 4 to 7 were used to normalize the body temperature of the fly to the left, while ellipses 1, 2, 3 and 8 were used to normalize the body temperature of the fly to the right.

We then averaged these temperatures over the three to five ellipses surrounding each fly. The average temperature of each fly was acquired by delimiting the body (abdomen + thorax) of the fly using the freehand tool included in the software FLIR ResearchIR Max.

More precisely, the normalizing procedure we used is the following. For every experiment *i*, the pair (data_norm)_i_ of temperature differences between the flies and the background on which they laid is given by

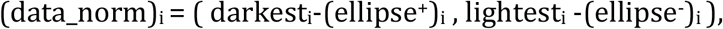

where

- darkest_i_ is the average over the first and second recordings in experiment *i* of the mean temperature measured on the abdomen and thorax of the darkest fly in the time interval [1 min, 2 min];
- lightest_i_ is the average over the first and second recordings in experiment *i* of the mean temperature of the lightest fly in the time interval [1 min, 2 min];
- (ellipse^+^)_i_ is the average over the two recordings and over the ellipses that are closest to the darkest fly in experiment *i* of the spatial and temporal mean temperature in each of these ellipses (the mean temperature of an ellipse being computed from a recording as the average over the time interval [1 min, 2 min] of the average over all pixels inside the ellipse of the temperature measured in these pixels).
- (ellipse^-^)_i_ is constructed in the same way as (ellipse^+^)_i_ but with the lightest fly.

The average over the first and second recordings of each pair of flies (after the positions of the flies were inverted) is taken in order to minimize the position effect. Moreover, the average temperature of the flies is computed over the time interval [1 min, 2 min] instead of the whole duration of the experiment [0 min, 3 min 30] to focus on the interval of time in which a relatively stable difference in temperatures between the two flies has established, after the initial increase and before switching off the lamp.

To test the hypothesis that the darkest fly becomes hotter that the lightest fly when both flies are exposed to light, we used a Wilcoxon signed rank test on the normalized paired measures of temperatures, for each of the following 4 groups of fly species or lines (with 13 to 15 pairs measured per group): *ebony^1^* and *yellow^1^ Drosophila melanogaster, Drosophila americana* and *Drosophila novamexicana*, *Drosophila yakuba* and *Drosophila santomea*, and *Drosophila melanogaster Dark* and *Pale* line*s*. The use of this statistical test requires that the distribution of the difference between the first and the second coordinate of (data_norm)_i_ (that is, the two standardized temperature measurements) within each group should be symmetrical about its mean. We confirmed that this assumption was indeed satisfied with another (one-dimensional) Wilcoxon signed rank test applied to the empirical distribution of these differences across replicate pairs within each group. The histograms obtained are displayed in Figure S1.

Based on the p-values for the Wilcoxon rank signed test, which are all larger than 0.05, the distribution of the histograms of [darkest_i_ - (ellipse^+^)_i_] - [lightest_i_ - (ellipse^-^)_i_] could be considered to be approximately symmetric about their means for the 4 groups of species or lines. This allowed us to perform Wilcoxon signed rank tests on paired samples for the 4 series of measurements of pairs of flies with different pigmentations.

### Measure of pigmentation differences

Photographs of flies were taken with a binocular equipped with Leica DC480 digital camera using *Leica IM50 Image Manager Software*. We took photos of pairs of flies corresponding to each of the four comparisons (10 pairs for each comparison). Using ImageJ, we decomposed each picture in hue, saturation and brightness and measured hue mean pixel intensity in thorax+abdomen of each fly. We then calculated the hue difference between the darkest and the lightest fly for each pair (Table S9-S12).

## Data availability statement

Raw data for temperatures and pigmentation are provided in supplementary information as tables S1-13.

## Acknowledgments

We thank Virginie Courtier-Orgogozo, the late Jean David and Jorge Vieira for drosophila stocks. We thank CNRS Mission for Transversal and Interdisciplinary Initiatives (MITI) for funding this research and LF PhD.

## Author contributions

Conception and design of the work: JMG; acquisition of data: LF and JMG; analysis and interpretation of data LF, JMG, AV, LMC, SM, MR and PC; drafting of the manuscript: LF, AV and JMG, with comments and suggestions from LMC, SM, MR and PC.

## Supplementary table and figure legends

Table S1: Data set for *D. melanogaster ebony* mutant females. The unit of measure is in Celsius and rounded down to nearest 0.01.

Table S2: Data set for *D. melanogaster yellow* mutant females. The unit of measure is in Celsius and rounded down to nearest 0.01.

Table S3: Data set for *D. americana* females. The unit of measure is in Celsius and rounded down to nearest 0.01.

Table S4: Data set for *D. novamexicana* females. The unit of measure is in Celsius and rounded down to nearest 0.01.

Table S5: Data set for *D.yakuba* males. The unit of measure is in Celsius and rounded down to nearest 0.01.

Table S6: Data set for *D. santomea* males. The unit of measure is in Celsius and rounded down to nearest 0.01.

Table S7: Data set for *D. melanogaster Dark* females. The unit of measure is in Celsius and rounded down to nearest 0.01.

Table S8: Data set for *D. melanogaster Pale* females. The unit of measure is in Celsius and rounded down to nearest 0.01.

Table S9: Data set for the differences in pigmentation (hue) between *D. melanogaster ebony* and *yellow* females

Table S10: Data set for the differences in pigmentation (hue) between *D. americana* and *D. novamexicana* females.

Table S11: Data set for differences in pigmentation (hue) between *D. yakuba* and *D. santomea* males.

Table S12: Data set for differences in pigmentation (hue) between *D. melanogaster Dark* and *Pale* females

Figure S1:

a: Histogram of the differences between the normalized temperature of the *D. melanogaster ebony* fly and that of the *D. melanogaster yellow* fly. It is set to have 7 bins. The box on the top-right of the picture is the result of a Wilcoxon signed rank exact test of the symmetry of the distribution of the difference between the two coordinates of data_norm with respect to its mean. The value of the test statistic is V = 57 and the associated p-value is 0.8871. Since this p-value is larger than 0.05, we cannot reject the hypothesis that the distribution of the temperature difference may be considered to be symmetric about its mean. The size of the data set is 15.

b: Histogram of the differences between the normalized temperature of *D. americana* and that of *D. novamexicana*. It is set to have 7 bins. The box on the top-right of the picture is the result of a Wilcoxon signed rank exact test to test the symmetry of the distribution of the difference between the two coordinates of data_norm with respect to its mean. The value of the test statistic is V = 43 and the associated p-value is 0.8926. Since the p-value is larger than 0.05, we cannot reject the hypothesis that the distribution of the temperature difference may be considered to be symmetric about its mean. The size of the data set is 13.

c: Histogram of the differences between the normalized temperature of *D. yakuba* and that of *D. santomea*. It is parametrised to have 7 bins. The box on the top-right of the picture is the result of a Wilcoxon signed rank exact test to test the symmetry of the distribution of the difference between the two coordinates of data_norm with respect to its mean. The value of the test statistic is V = 43 and the associated p-value is 0.8926. Since the p-value is larger than 0.05, we cannot reject the hypothesis that the distribution of the temperature difference may be considered to be symmetric about its mean. The size of the data set is 13.

d: Histogram of the differences between the normalized temperature of *D. melanogaster Dark* line and that of *D. melanogaster Pale* line. It is set to have 7 bins. The box on the top-right of the picture is the result of a Wilcoxon signed rank exact test to test the symmetry of the distribution of the difference between the two coordinates of data_norm with respect to its mean. The test statistic is V = 42 and the associated p- value is 0.8394. Here again, we cannot reject the hypothesis that the distribution of the temperature difference may be considered to be symmetric about its mean. The size of the data set is 13.

