## Supplementary tables and figure for "Drosophilid cuticle pigmentation impacts body temperature": Table S1.docx

|  | **Before inversion** | | | **After inversion** | | | **Mean between two inversions** | | |
| --- | --- | --- | --- | --- | --- | --- | --- | --- | --- |
| In ° | Fly | Ellipse | Fly - Ellipse | Fly | Ellipse | Fly - Ellipse | Fly | Ellipse | Fly - Ellipse |
| **1** | 22.30 | 20.13 | 2.17 | 21.92 | 19.75 | 2.17 | 22.11 | 19.94 | 2.17 |
| **2** | 22.31 | 20.11 | 2.20 | 22.40 | 20.18 | 2.22 | 22.35 | 20.14 | 2.21 |
| **3** | 22.49 | 20.48 | 2.01 | 22.28 | 20.24 | 2.04 | 22.38 | 20.36 | 2.02 |
| **4** | 22.57 | 20.18 | 2.39 | 22.75 | 20.19 | 2.56 | 22.66 | 20.18 | 2.48 |
| **5** | 22.12 | 20.06 | 2.06 | 22.29 | 20.23 | 2.06 | 22.20 | 20.14 | 2.06 |
| **6** | 22.18 | 20.06 | 2.12 | 22.55 | 20.08 | 2.47 | 22.36 | 20.07 | 2.29 |
| **7** | 22.15 | 20.01 | 2.14 | 22.22 | 20.21 | 2.01 | 22.18 | 20.11 | 2.07 |
| **8** | 22.62 | 20.33 | 2.29 | 22.62 | 20.25 | 2.37 | 22.62 | 20.29 | 2.33 |
| **9** | 21.84 | 19.69 | 2.15 | 21.98 | 19.84 | 2.14 | 21.91 | 19.76 | 2.15 |
| **10** | 22.38 | 20.23 | 2.15 | 21.68 | 19.82 | 1.86 | 22.03 | 20.02 | 2.01 |
| **11** | 22.84 | 20.44 | 2.40 | 22.20 | 20.13 | 2.07 | 22.52 | 20.28 | 2.24 |
| **12** | 23.03 | 20.43 | 2.60 | 22.63 | 20.16 | 2.47 | 22.83 | 20.29 | 2.54 |
| **13** | 22.54 | 19.89 | 2.65 | 22.69 | 20.00 | 2.69 | 22.61 | 19.94 | 2.67 |
| **14** | 20.65 | 18.86 | 1.79 | 21.67 | 19.47 | 2.20 | 21.16 | 19.16 | 2.00 |
| **15** | 23.41 | 20.40 | 2.99 | 22.99 | 19.80 | 3.19 | 23.20 | 20.10 | 3.10 |
