## Supplementary tables and figure for "Drosophilid cuticle pigmentation impacts body temperature": Table S2.docx

|  | **Before inversion** | | | **After inversion** | | | **Mean between two inversions** | | |
| --- | --- | --- | --- | --- | --- | --- | --- | --- | --- |
| In ° | Fly | Ellipse | Fly - Ellipse | Fly | Ellipse | Fly - Ellipse | Fly | Ellipse | Fly - Ellipse |
| **1** | 21.42 | 19.87 | 1.55 | 21.23 | 19.87 | 1.36 | 21.32 | 19.87 | 1.45 |
| **2** | 21.50 | 19.93 | 1.57 | 21.85 | 20.17 | 1.68 | 21.67 | 20.05 | 1.62 |
| **3** | 21.89 | 20.26 | 1.63 | 22.10 | 20.25 | 1.85 | 21.99 | 20.25 | 1.74 |
| **4** | 21.80 | 20.01 | 1.79 | 22.04 | 20.18 | 1.86 | 21.92 | 20.09 | 1.83 |
| **5** | 21.38 | 19.83 | 1.55 | 21.69 | 20.04 | 1.65 | 21.53 | 19.93 | 1.60 |
| **6** | 21.26 | 19.94 | 1.32 | 21.82 | 20.20 | 1.62 | 21.54 | 20.07 | 1.47 |
| **7** | 21.68 | 20.14 | 1.54 | 21.63 | 19.93 | 1.70 | 21.65 | 20.03 | 1.62 |
| **8** | 21.85 | 20.08 | 1.77 | 22.07 | 20.30 | 1.77 | 21.96 | 20.19 | 1.77 |
| **9** | 21.37 | 19.73 | 1.64 | 21.24 | 19.71 | 1.53 | 21.30 | 19.72 | 1.58 |
| **10** | 21.79 | 20.42 | 1.37 | 21.32 | 19.70 | 1.62 | 21.55 | 20.06 | 1.49 |
| **11** | 21.93 | 20.23 | 1.70 | 21.49 | 20.12 | 1.37 | 21.71 | 20.17 | 1.54 |
| **12** | 22.05 | 20.28 | 1.77 | 22.07 | 20.06 | 2.01 | 22.06 | 20.17 | 1.89 |
| **13** | 21.37 | 19.75 | 1.62 | 21.60 | 19.93 | 1.67 | 21.48 | 19.84 | 1.64 |
| **14** | 19.98 | 18.66 | 1.32 | 20.93 | 19.50 | 1.43 | 20.45 | 19.08 | 1.37 |
| **15** | 22.55 | 20.21 | 2.34 | 21.89 | 19.88 | 2.01 | 22.22 | 20.04 | 2.18 |
