## Supplementary tables and figure for "Drosophilid cuticle pigmentation impacts body temperature": Table S3.docx

|  | **Before inversion** | | | **After inversion** | | | **Mean between two inversions** | | |
| --- | --- | --- | --- | --- | --- | --- | --- | --- | --- |
| In ° | Fly | Ellipse | Fly - Ellipse | Fly | Ellipse | Fly - Ellipse | Fly | Ellipse | Fly - Ellipse |
| **1** | 23.71 | 20.09 | 3.62 | 23.83 | 20.00 | 3.83 | 23.77 | 20.04 | 3.72 |
| **2** | 23.61 | 20.72 | 2.89 | 23.09 | 20.26 | 2.83 | 23.35 | 20.49 | 2.86 |
| **3** | 24.14 | 20.39 | 3.75 | 23.61 | 20.25 | 3.36 | 23.87 | 20.32 | 3.55 |
| **4** | 23.97 | 20.45 | 3.52 | 23.19 | 20.26 | 2.93 | 23.58 | 20.35 | 3.22 |
| **5** | 24.42 | 20.53 | 3.89 | 24.45 | 20.55 | 3.90 | 24.43 | 20.54 | 3.89 |
| **6** | 24.42 | 20.88 | 3.54 | 24.46 | 21.06 | 3.40 | 24.44 | 20.97 | 3.47 |
| **7** | 23.57 | 20.34 | 3.23 | 24.54 | 21.13 | 3.41 | 24.05 | 20.73 | 3.32 |
| **8** | 23.32 | 19.79 | 3.53 | 23.82 | 20.79 | 3.03 | 23.57 | 20.29 | 3.28 |
| **9** | 23.02 | 19.96 | 3.06 | 22.92 | 19.97 | 2.95 | 22.97 | 19.96 | 3.00 |
| **10** | 23.45 | 20.27 | 3.18 | 23.28 | 20.28 | 3.00 | 23.36 | 20.27 | 3.09 |
| **11** | 22.82 | 20.18 | 2.64 | 22.67 | 19.95 | 2.72 | 22.74 | 20.06 | 2.68 |
| **12** | 23.12 | 20.12 | 3.00 | 22.82 | 19.93 | 2.89 | 22.97 | 20.02 | 2.94 |
| **13** | 23.30 | 20.12 | 3.18 | 22.90 | 20.01 | 2.89 | 23.10 | 20.06 | 3.03 |
