## Supplementary tables and figure for "Drosophilid cuticle pigmentation impacts body temperature": Table S4.docx

|  | **Before inversion** | | | **After inversion** | | | **Mean between two inversions** | | |
| --- | --- | --- | --- | --- | --- | --- | --- | --- | --- |
| In ° | Fly | Ellipse | Fly - Ellipse | Fly | Ellipse | Fly - Ellipse | Fly | Ellipse | Fly - Ellipse |
| **1** | 22.97 | 20.01 | 2.96 | 23.09 | 20.04 | 3.05 | 23.03 | 20.02 | 3.00 |
| **2** | 23.40 | 20.93 | 2.47 | 22.62 | 19.86 | 2.76 | 23.01 | 20.39 | 2.61 |
| **3** | 22.95 | 20.36 | 2.59 | 22.55 | 19.96 | 2.59 | 22.75 | 20.16 | 2.59 |
| **4** | 23.35 | 20.84 | 2.51 | 22.78 | 19.90 | 2.88 | 23.06 | 20.37 | 2.69 |
| **5** | 23.40 | 20.63 | 2.77 | 23.18 | 20.15 | 3.03 | 23.29 | 20.39 | 2.90 |
| **6** | 24.39 | 21.48 | 2.91 | 23.73 | 20.98 | 2.75 | 24.06 | 21.23 | 2.83 |
| **7** | 22.75 | 20.24 | 2.51 | 23.68 | 20.99 | 2.69 | 23.21 | 20.61 | 2.60 |
| **8** | 22.70 | 19.92 | 2.78 | 23.07 | 20.35 | 2.72 | 22.88 | 20.13 | 2.75 |
| **9** | 22.62 | 19.83 | 2.79 | 22.24 | 19.80 | 2.44 | 22.43 | 19.81 | 2.61 |
| **10** | 22.70 | 20.44 | 2.26 | 22.99 | 20.30 | 2.69 | 22.84 | 20.37 | 2.47 |
| **11** | 22.32 | 20.23 | 2.09 | 22.14 | 19.87 | 2.27 | 22.23 | 20.05 | 2.18 |
| **12** | 22.43 | 20.14 | 2.29 | 22.21 | 19.93 | 2.28 | 22.32 | 20.03 | 2.28 |
| **13** | 22.74 | 20.20 | 2.54 | 22.55 | 19.94 | 2.61 | 22.64 | 20.07 | 2.57 |
