## Supplementary tables and figure for "Drosophilid cuticle pigmentation impacts body temperature": Table S5.docx

|  | **Before inversion** | | | **After inversion** | | | **Mean between two inversions** | | |
| --- | --- | --- | --- | --- | --- | --- | --- | --- | --- |
| In ° | Fly | Ellipse | Fly - Ellipse | Fly | Ellipse | Fly - Ellipse | Fly | Ellipse | Fly - Ellipse |
| **1** | 20.76 | 19.27 | 1.49 | 20.75 | 19.08 | 1.67 | 20.75 | 19.17 | 1.58 |
| **2** | 20.61 | 19.08 | 1.53 | 20.86 | 19.14 | 1.72 | 20.73 | 19.11 | 1.62 |
| **3** | 20.72 | 19.25 | 1.47 | 20.80 | 19.22 | 1.58 | 20.76 | 19.23 | 1.52 |
| **4** | 20.87 | 19.03 | 1.84 | 20.86 | 19.05 | 1.81 | 20.86 | 19.04 | 1.82 |
| **5** | 20.83 | 19.30 | 1.53 | 20.43 | 19.02 | 1.41 | 20.63 | 19.16 | 1.47 |
| **6** | 20.61 | 19.19 | 1.42 | 20.75 | 18.97 | 1.78 | 20.68 | 19.08 | 1.60 |
| **7** | 20.70 | 19.39 | 1.31 | 20.85 | 19.40 | 1.45 | 20.77 | 19.39 | 1.38 |
| **8** | 20.55 | 19.37 | 1.18 | 20.49 | 19.26 | 1.23 | 20.52 | 19.31 | 1.20 |
| **9** | 20.73 | 19.36 | 1.37 | 21.03 | 19.40 | 1.63 | 20.88 | 19.38 | 1.50 |
| **10** | 21.89 | 19.78 | 2.11 | 21.38 | 19.72 | 1.66 | 21.63 | 19.75 | 1.88 |
| **11** | 21.69 | 19.93 | 1.76 | 21.74 | 19.63 | 2.11 | 21.71 | 19.78 | 1.93 |
| **12** | 21.92 | 20.12 | 1.80 | 21.80 | 20.09 | 1.71 | 21.86 | 20.10 | 1.75 |
| **13** | 21.74 | 20.11 | 1.63 | 21.79 | 19.93 | 1.86 | 21.76 | 20.02 | 1.74 |
