## Supplementary tables and figure for "Drosophilid cuticle pigmentation impacts body temperature": Table S6.docx

|  | **Before inversion** | | | **After inversion** | | | **Mean between two inversions** | | |
| --- | --- | --- | --- | --- | --- | --- | --- | --- | --- |
| In ° | Fly | Ellipse | Fly - Ellipse | Fly | Ellipse | Fly - Ellipse | Fly | Ellipse | Fly - Ellipse |
| **1** | 20.64 | 19.17 | 1.47 | 20.57 | 19.18 | 1.39 | 20.60 | 19.17 | 1.43 |
| **2** | 20.43 | 19.01 | 1.42 | 20.74 | 19.12 | 1.62 | 20.58 | 19.06 | 1.52 |
| **3** | 20.65 | 19.21 | 1.44 | 20.86 | 19.26 | 1.60 | 20.75 | 19.23 | 1.52 |
| **4** | 20.73 | 18.94 | 1.79 | 20.78 | 19.15 | 1.63 | 20.75 | 19.04 | 1.71 |
| **5** | 20.81 | 19.19 | 1.62 | 20.20 | 19.00 | 1.20 | 20.50 | 19.09 | 1.41 |
| **6** | 20.47 | 19.11 | 1.36 | 20.50 | 19.05 | 1.45 | 20.48 | 19.08 | 1.40 |
| **7** | 20.76 | 19.23 | 1.53 | 20.80 | 19.45 | 1.35 | 20.78 | 19.34 | 1.44 |
| **8** | 20.33 | 19.25 | 1.08 | 20.13 | 19.21 | 0.92 | 20.23 | 19.23 | 1.00 |
| **9** | 20.41 | 19.29 | 1.12 | 20.40 | 19.43 | 0.97 | 20.40 | 19.36 | 1.04 |
| **10** | 21.47 | 19.63 | 1.84 | 21.47 | 19.84 | 1.63 | 21.47 | 19.73 | 1.73 |
| **11** | 21.31 | 19.67 | 1.64 | 21.53 | 19.87 | 1.66 | 21.42 | 19.77 | 1.65 |
| **12** | 21.68 | 19.89 | 1.79 | 21.80 | 20.23 | 1.57 | 21.74 | 20.06 | 1.68 |
| **13** | 21.60 | 19.88 | 1.72 | 21.60 | 20.13 | 1.47 | 21.60 | 20.00 | 1.59 |
