## Supplementary tables and figure for "Drosophilid cuticle pigmentation impacts body temperature": Table S7.docx

|  | **Before inversion** | | | **After inversion** | | | **Mean between two inversions** | | |
| --- | --- | --- | --- | --- | --- | --- | --- | --- | --- |
| In ° | Fly | Ellipse | Fly - Ellipse | Fly | Ellipse | Fly - Ellipse | Fly | Ellipse | Fly - Ellipse |
| **1** | 22.37 | 20.51 | 1.86 | 22.11 | 20.20 | 1.91 | 22.24 | 20.35 | 1.88 |
| **2** | 21.96 | 20.18 | 1.78 | 21.71 | 19.85 | 1.86 | 21.83 | 20.01 | 1.82 |
| **3** | 22.10 | 19.95 | 2.15 | 22.35 | 20.52 | 1.83 | 22.22 | 20.23 | 1.99 |
| **4** | 22.10 | 19.88 | 2.22 | 22.16 | 19.96 | 2.20 | 22.13 | 19.92 | 2.21 |
| **5** | 21.62 | 20.21 | 1.41 | 20.97 | 19.28 | 1.69 | 21.29 | 19.74 | 1.55 |
| **6** | 21.50 | 19.89 | 1.61 | 21.77 | 19.91 | 1.86 | 21.63 | 19.90 | 1.73 |
| **7** | 21.14 | 19.30 | 1.84 | 22.14 | 20.31 | 1.83 | 21.64 | 19.80 | 1.83 |
| **8** | 21.93 | 19.94 | 1.99 | 21.82 | 20.00 | 1.82 | 21.87 | 19.97 | 1.90 |
| **9** | 22.01 | 20.25 | 1.76 | 21.26 | 19.37 | 1.89 | 21.63 | 19.81 | 1.82 |
| **10** | 21.65 | 20.00 | 1.65 | 21.41 | 19.66 | 1.75 | 21.53 | 19.83 | 1.70 |
| **11** | 21.55 | 19.74 | 1.81 | 21.94 | 20.18 | 1.76 | 21.74 | 19.96 | 1.78 |
| **12** | 22.05 | 20.15 | 1.90 | 21.34 | 19.45 | 1.89 | 21.69 | 19.80 | 1.89 |
| **13** | 21.84 | 20.18 | 1.66 | 21.62 | 19.53 | 2.09 | 21.73 | 19.85 | 1.87 |
