## Supplementary tables and figure for "Drosophilid cuticle pigmentation impacts body temperature": Table S8.docx

|  | **Before inversion** | | | **After inversion** | | | **Mean between two inversions** | | |
| --- | --- | --- | --- | --- | --- | --- | --- | --- | --- |
| In ° | Fly | Ellipse | Fly - Ellipse | Fly | Ellipse | Fly - Ellipse | Fly | Ellipse | Fly - Ellipse |
| **1** | 22.38 | 20.66 | 1.72 | 21.82 | 20.09 | 1.73 | 22.10 | 20.37 | 1.72 |
| **2** | 22.05 | 20.33 | 1.72 | 21.51 | 19.74 | 1.77 | 21.78 | 20.03 | 1.74 |
| **3** | 21.77 | 20.14 | 1.63 | 22.02 | 20.24 | 1.78 | 21.89 | 20.19 | 1.70 |
| **4** | 21.70 | 19.24 | 2.46 | 21.47 | 20.17 | 1.30 | 21.58 | 19.70 | 1.88 |
| **5** | 21.62 | 20.13 | 1.49 | 21.13 | 19.41 | 1.72 | 21.37 | 19.77 | 1.60 |
| **6** | 21.36 | 19.83 | 1.53 | 21.82 | 20.05 | 1.77 | 21.59 | 19.94 | 1.65 |
| **7** | 20.83 | 19.15 | 1.68 | 22.10 | 20.29 | 1.81 | 21.46 | 19.72 | 1.74 |
| **8** | 21.64 | 19.83 | 1.81 | 21.73 | 20.14 | 1.59 | 21.68 | 19.98 | 1.70 |
| **9** | 21.79 | 20.13 | 1.66 | 21.28 | 19.52 | 1.76 | 21.53 | 19.82 | 1.71 |
| **10** | 21.33 | 19.99 | 1.34 | 21.21 | 19.69 | 1.52 | 21.27 | 19.84 | 1.43 |
| **11** | 21.29 | 19.59 | 1.70 | 21.96 | 20.26 | 1.70 | 21.62 | 19.92 | 1.70 |
| **12** | 21.73 | 20.10 | 1.63 | 21.54 | 19.59 | 1.95 | 21.63 | 19.84 | 1.79 |
| **13** | 21.73 | 20.03 | 1.70 | 21.66 | 19.74 | 1.92 | 21.69 | 19.88 | 1.81 |
