## Supplementary figures and images for "Drosophilid cuticle pigmentation impacts body temperature"

### Figure S1.tif

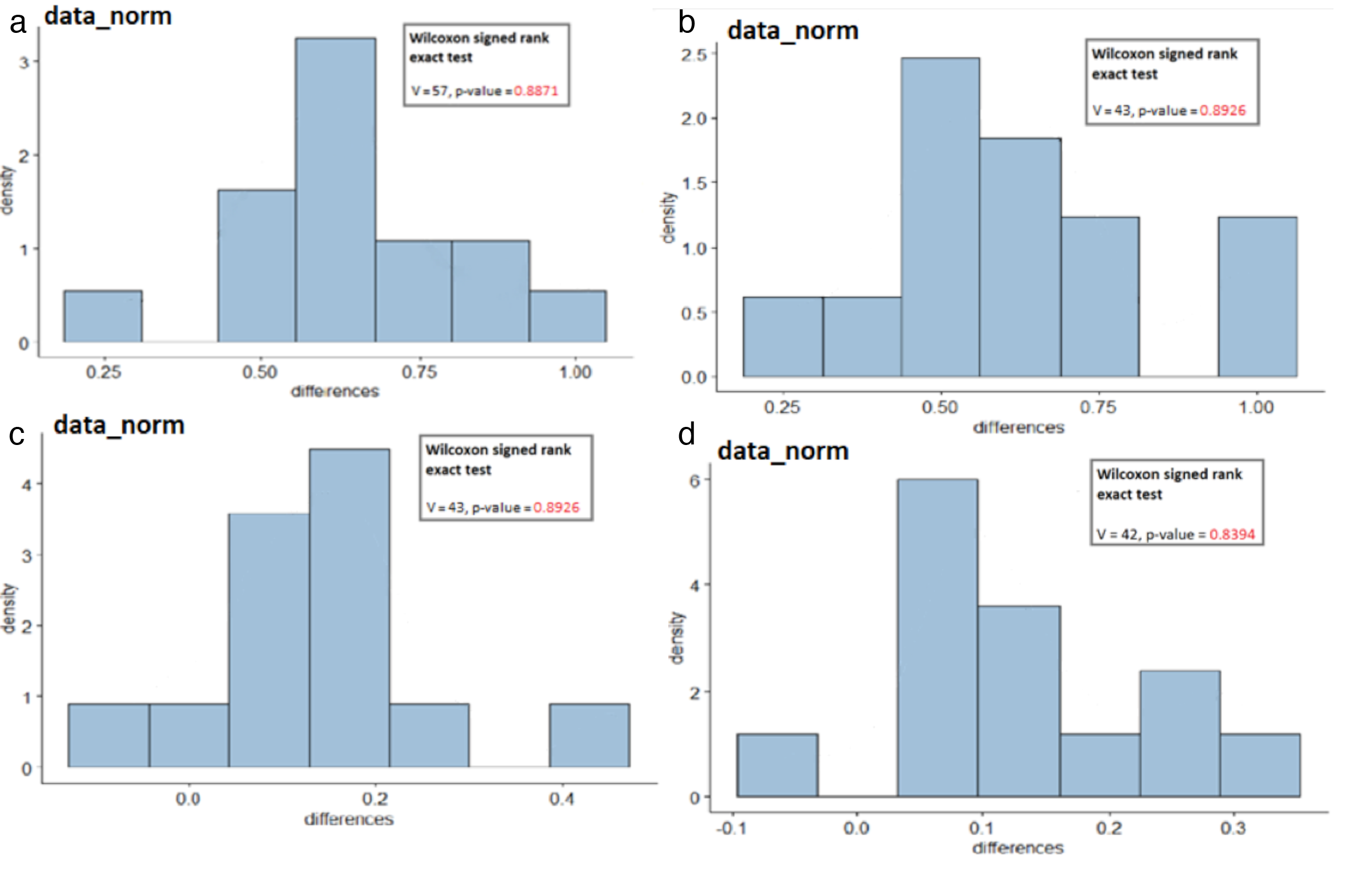
